# Neural Flip-Flops III: Stomatogastric Ganglion

**DOI:** 10.1101/2020.11.29.403154

**Authors:** Lane Yoder

**Affiliations:** Department of Science and Mathematics, retired, University of Hawaii, Kapiolani, Honolulu, Hawaii

**Keywords:** stomatogastric ganglion, STG, central pattern generator, CPG, pylorus, flip-flop, JK flip-flop, oscillator, toggle, bursting neuron, explicit neural model, neuronal network

## Abstract

The stomatogastric ganglion (STG) is a group of about 30 neurons that resides on the stomach in decapod crustaceans. Its two central pattern generators (CPGs) control the chewing action of the gastric mill and the peristaltic movement of food through the pylorus to the gut. The STG has been studied extensively because it has properties that are common to all nervous systems and because of the small number of neurons and other features that make it convenient to study. So many details are known that the STG is considered a classic test case in neuroscience for the reductionist strategy of explaining the emergence of macro-level phenomena from micro-level data. In spite of the intense scrutiny the STG has received, how it generates its rhythmic patterns of bursts remains unknown.

The explicit neural networks proposed here model the pyloric CPG of the American lobster (*Homarus americanus*). The models share enough significant features with the lobster’s CPG that they may be considered first approximations, or perhaps simplified versions, of STG architecture. The similarities include 1) mostly inhibitory synapses; 2) pairs of cells with reciprocal inhibitory inputs, complementary outputs that are approximately 180 degrees out of phase, and state changes occurring with the high output changing first; 3) cells that have reciprocal, inhibitory inputs with more than one other cell; and 4) six cells that produce coordinated oscillations with the same period, four phases distributed approximately uniformly over the period, and half of the burst durations approximately 1/4 of the period and the other half 3/8.

Each model’s connectivity is explicit, and its operation depends only on minimal neuron capabilities of excitation and inhibition. One model performs a function that fills a gap in standard ring oscillators. It is apparently new to engineering, making it an example of neuroscience and logic circuit design informing each other.

Some models are derived from standard circuit designs by moving each negation symbol from one end of a connection to the other. This does not change the logic of the network, but it changes each logic gate to one that can be implemented with a single neuron.

## 1. Introduction

This article is the sixth in a series of articles that show how neurons are likely to be connected to perform certain brain functions. The first three articles [1-3] showed how a fuzzy logic decoder can generate the phenomena central to color vision and olfaction and can account for major aspects of the anatomical structure and physiological organization of the neocortex. The next two articles [4, 5] showed how Boolean logic neural flip-flops (NFFs) and NFFs configured as central pattern generators (CPGs) can produce detailed characteristics of short-term memory and electroencephalography.

The present article presents several novel CPG designs that show how neurons can be connected to produce rhythmic bursts that closely match the oscillations produced by the American lobster’s pyloric CPG. These logic circuits are some of the simplest oscillators that can be constructed with components whose only active properties are excitation and inhibition. All of them produce oscillations that are close to the pyloric CPG in phases and burst durations. This supports the hypothesis that the pyloric CPG is fundamentally similar in design to one or more of the models. None of the models is an exact match, and each is close to the pyloric in different ways. The additional complexity of the pyloric CPG apparently alters the signals slightly.

The novel flip-flop ring oscillator proposed here is a generalization of the standard inverter ring oscillator, which is composed of an odd number of inverters connected in a ring. A flip-flop ring is composed of two or more of the simplest flip-flops, with both outputs of each flip-flop connected as inputs to the next one. Flip-flop rings fill a gap in the patterns produced by inverter rings. The two ring types produce, respectively, even and odd numbers of oscillations, with their phases distributed approximately uniformly over one cycle. An inverter ring could not produce the four pyloric oscillations of the American lobster.

The model CPGs can be implemented with neurons or electronic hardware. The models were simulated with minimal properties of neuron properties of excitation and inhibition and with electronic circuit simulation software. The neural and electronic simulations returned virtually identical results. The close fits between the electronic simulations and pyloric oscillations are a remarkable convergence of phenomena generated by any two different mechanisms, but especially for mechanisms that are so different in materials (electronic hardware versus lobster neurons), network complexity (16 or 24 transistors, connected in simple, repeating patterns, versus the more complex pyloric CPG), design methods (modern engineering versus natural selection) and speed (the electronic and lobster frequencies differ by a factor of about 17 million). The similar results, along with the similar operational properties of the components and similar connectivity between components, suggest the results are produced in the same way, at least in principle if not in the details.

A few other concepts presented here are also likely to be new:

Neurons are well known to have a thousand or more connections. It is shown here how complex logic circuits can be implemented with a few neurons and many synapses.

Some of the neural networks presented here can be derived from standard logic circuit designs by the simple transformation of moving each negation symbol from one end of a connection to the other. Many more networks that can be implemented with neurons can be derived by this simple method and may explain more nervous system functions. Although the modifications are minor, both the method and the derived networks are likely to be new to engineering, as well as to neuroscience.

The observation that the burst centers of four of the American lobster’s pyloric oscillations are approximately uniformly distributed over the period is apparently new. The different models presented here show that a CPG can produce oscillations with phases uniformly distributed at the burst centers, at the burst onsets, or both.

Two of the model CPGs demonstrate how a neuron (or an electronic logic gate) can produce useful functions with reciprocal, inhibitory inputs with more than one other neuron. The STG has several such neurons. Although pairs of logic gates with reciprocal, inhibitory inputs are quite common in electronic logic circuits, electronic gates apparently do not have reciprocal, inhibitory inputs with more than one other gate.

Several testable predictions are given here, and STG phenomena are shown to support several predictions of neural flip-flops that were given in a previous paper on short-term memory.

## 2. Methods

### 2.1. Neural Boolean logic

All Boolean logic results for the networks presented here follow from the neuron responses to binary (high and low) input signals, given in two tables below, and the algebra of Boolean logic applied to the networks’ connections. Analog signals (intermediate strengths between high and low) were considered in [4, 5] only to show how NFFs can generate robust binary signals in the presence of moderate levels of additive noise in binary inputs. That discussion will not be repeated here.

#### 2.1.1. Binary neuron signals

Neuron signal strength, or intensity, is normalized here by dividing it by the maximum possible intensity for the given level of adaptation. This puts intensities in the interval from 0 to 1, with 0 meaning no signal and 1 meaning the maximum intensity. The normalized number is called the *response intensity* or simply the *response* of the neuron. Normalization is only for convenience. Non-normalized signal strengths, with the highest and lowest values labeled Max & Min rather than 1 and 0, would do as well.

The responses 1 and 0 are called binary signals collectively and high and low signals separately. If 1 and 0 stand for the Boolean truth values TRUE and FALSE, neurons can process information contained in binary signals by functioning as logic operators.

The strength of a signal consisting of action potentials, or spikes, can be measured by spike frequency. A high signal consists of a burst of spikes at the maximum spiking rate. For a signal that oscillates between high and low, the frequency of the oscillation is the frequency of bursts, not the frequency of spikes.

For binary signals, the response of a neuron with one excitatory and one inhibitory input is assumed to be as shown in Table 1. Of the 16 possible binary functions of two variables, this table represents the only one that is consistent with the customary meanings of “excitation” and “inhibition.” Table 1 is also a logic truth table, with the last column representing the truth values of the statement X AND NOT Y. In simplest terms, the neuron performs this logic function because it is active when it has excitatory input *and* does *not* have inhibitory input.

**Table 1.**
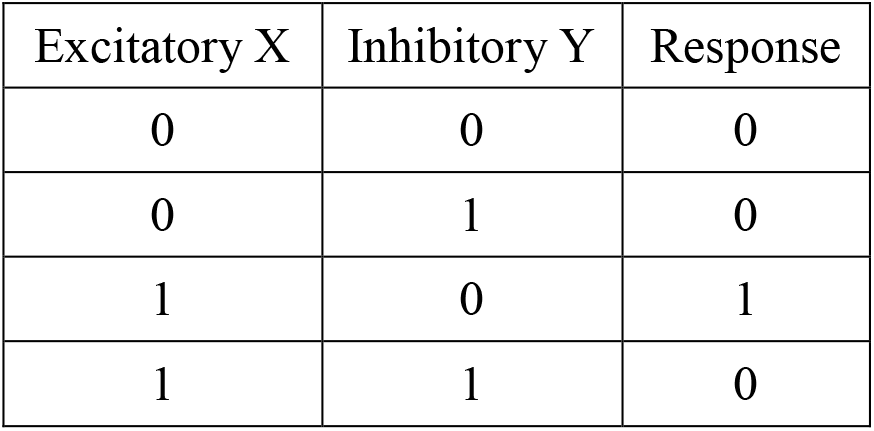
Neuron response to two binary inputs, one excitatory and one inhibitory. The response is the logical truth value of X AND NOT Y.

| Excitatory X | Inhibitory Y | Response |
| --- | --- | --- |
| 0 | 0 | 0 |
| 0 | 1 | 0 |
| 1 | 0 | 1 |
| 1 | 1 | 0 |

The AND-NOT gate is virtually never used as a building block in logic circuit design. Its significance for the networks presented here is that it can be implemented with a single neuron. It was shown in [2, 4] that the AND-NOT gate, with access to high input, is functionally complete, meaning any logic function can be performed by a network of such components.

Some neurons are active spontaneously and continuously without excitatory input [6,7]. In the figures, neurons with no excitatory input are spontaneously active. The behavior of a spontaneously active neuron is assumed to be the same as a neuron with a high excitatory input.

#### 2.1.2. Single neuron logic primitives

Fig 1 shows symbols for a few logic primitives. For several reasons that were detailed in [4], networks are illustrated with standard (ANSI/IEEE) logic symbols rather than symbols commonly used in neuroscience schematic diagrams. One of the reasons is that the symbols can be interpreted in two ways. As a logic symbol, the rectangle with one rounded side in Fig 1A represents the AND logic function, and the circle represents negation. The input variables X and Y represent truth values TRUE or FALSE, and the output represents the truth value X AND NOT Y. Second, Fig 1A can also represent a single neuron, with a circle representing inhibitory input and no circle representing excitatory input. If X and Y are binary inputs, the output is X AND NOT Y by Table 1. For the rest of the symbols in Fig 1 and the networks in subsequent figures, the outputs follow from straightforward Boolean logic.

**Fig 1.**
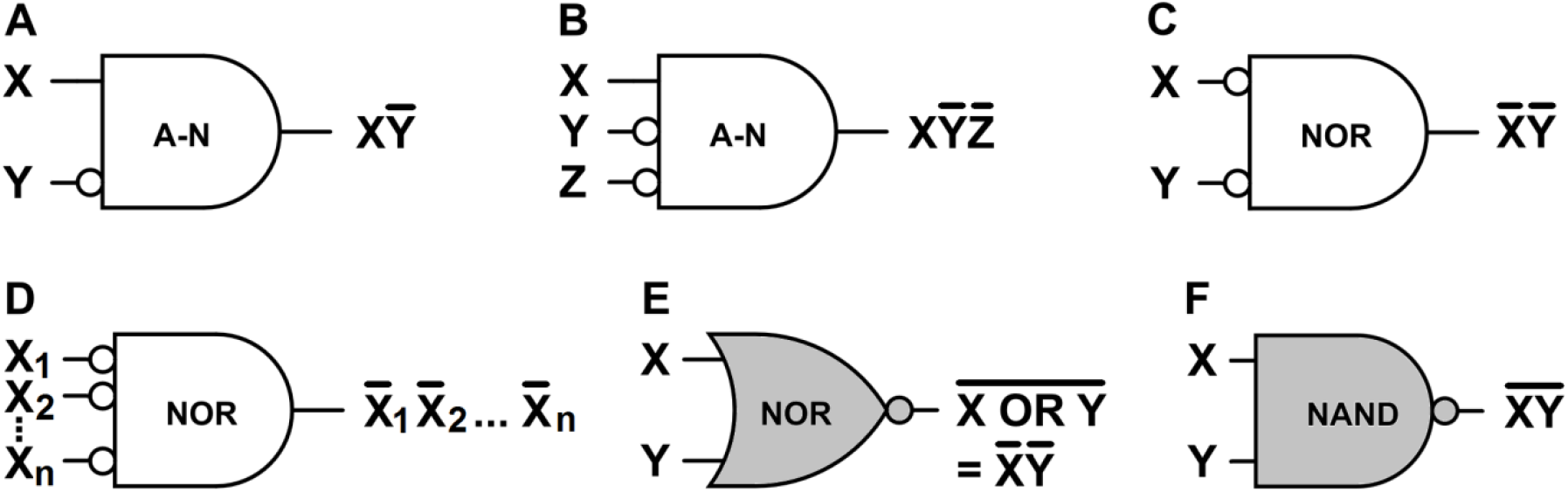
Logic primitives: AND-NOT, NOR, NAND. Each white diagram can be implemented with a single neuron or with electronic components. The gray symbols are commonly used in electronic logic circuit design. **A**. A logic symbol for an AND-NOT gate, or a neuron with one excitatory input and one inhibitory input. **B**. A three-input AND-NOT gate, or a neuron with one excitatory input and two inhibitory inputs. **C**. The NOR function implemented with an AND gate or with a spontaneously active neuron and two inhibitory inputs. **D**. A multi-input NOR gate (short for “NOT OR”), or a spontaneously active neuron with several inhibitory inputs. **E**. The most common symbol for a NOR gate. **F**. The most common symbol for a NAND gate (for “NOT AND”).

The neuron logic for Fig 1B follows from Table 1: if one inhibitory input can suppress one excitatory input, then either one of two inhibitory inputs can suppress the excitatory input. The neuron output for Fig 1C follows from Fig 1B and the property that the behavior of a spontaneously active neuron is the same as a neuron with high excitatory input.

The logic primitive NOT(X OR Y) is called the NOR operator (for “NOT OR”). By De Morgan’s law, NOT(X OR Y) is logically equivalent to (NOT X) AND (NOT Y). The latter is the output of Fig 1C. In simplest terms, this neuron is a NOR gate because it is active when it has inhibitory input from neither X *nor* Y. Like the AND-NOT operator, the NOR operator is functionally complete. This operator will be used extensively in the figures and simulations, so its response function is shown in Table 2.

**Table 2.** A spontaneously active neuron’s NOR response to binary, inhibitory inputs (Fig 1C). The response (NOT X) AND (NOT Y) is logically equivalent to NOT(X OR Y) by de Morgan’s law.

| Inhibitory X | Inhibitory Y | Response |
| --- | --- | --- |
| 0 | 0 | 1 |
| 0 | 1 | 0 |
| 1 | 0 | 0 |
| 1 | 1 | 0 |

Fig 1D generalizes Fig 1C: Any of several inhibitory inputs can suppress a spontaneously active neuron. This general NOR gate is an efficient and powerful mechanism. It was shown in [2] that a neuron with a continuously high excitatory input and an inhibitory input X functions as a NOT gate (output NOT X), also known as an inverter. Negating the output of Fig 1D with an inverter makes the two-neuron network a general OR gate (output X_1_ OR X_2_ OR … OR X_n_) by De Morgan’s law. Negating some or all of the inputs to Fig 1D with inverters makes the NOR gate a general AND gate. This means any conjunction of n inputs or their negations can be implemented with at most n + 1 neurons and as few as 1, depending on how many of the inputs need to be negated. A well-known theorem of logic says that any logic statement can be expressed either as a sum (OR) of products (AND) or as a product of sums. This implies that complex logic can be implemented with a few neurons and many synapses.

Fig 1E is the most commonly used symbol for a NOR gate. The logic operations performed by Figs 1C and 1E are logically equivalent by de Morgan’s law. The symbol in Fig 1C will be used to represent the NOR gate in the network diagrams here because it is more easily interpreted as a single neuron with the inhibitory inputs indicated explicitly, and because it is convenient for showing how some neural networks can be derived from standard electronic designs.

The logic primitive NOT(X AND Y) is called the NAND operator (for “NOT AND”). Fig 1F is the most commonly used symbol for a NAND gate. The NAND operator is functionally complete, and the NAND and NOR gates are two of the most commonly used logic gates in electronic logic circuit design. Here the NAND gate will be used only in demonstrating the derivation of the neural CPGs from electronic circuits.

#### 2.1.3. Flip-flops and oscillators

An oscillator is the basic element of a timing mechanism. It produces periodic bursts of a high signal followed a quiescent interval of a low signal.

A flip-flop is a memory mechanism that stores one bit of information in an output that is either 0 or 1. This output is the flip-flop *state* or *memory bit*. A change in the state *inverts* the state. The information is stored by means of a brief input signal that determines the output. A distinction is sometimes made between a “flip-flop” and a “latch,” with the latter term reserved for asynchronous memory mechanisms that are not controlled by an oscillator. The more familiar “flip-flop” will be used here for all cases.

A toggle is a flip-flop with one input that inverts the flip-flop’s memory state each time the input is high. With input from an oscillator, a toggle functions as another oscillator. Because two high inputs are required for each cycle, a toggle-as-oscillator produces a signal whose frequency is exactly half that of the toggle’s input.

##### 2.1.3.1. Neural and electronic flip-flops and oscillators

Fig 2 shows several examples of flip-flops and oscillators. The flip-flops’ memory bit is labeled M in the diagrams. Input S *sets* the state to M = 1, and R *resets* it to M = 0. Feedback maintains a stable state. Configured as a toggle, the single input is labeled T.

**Fig 2.**
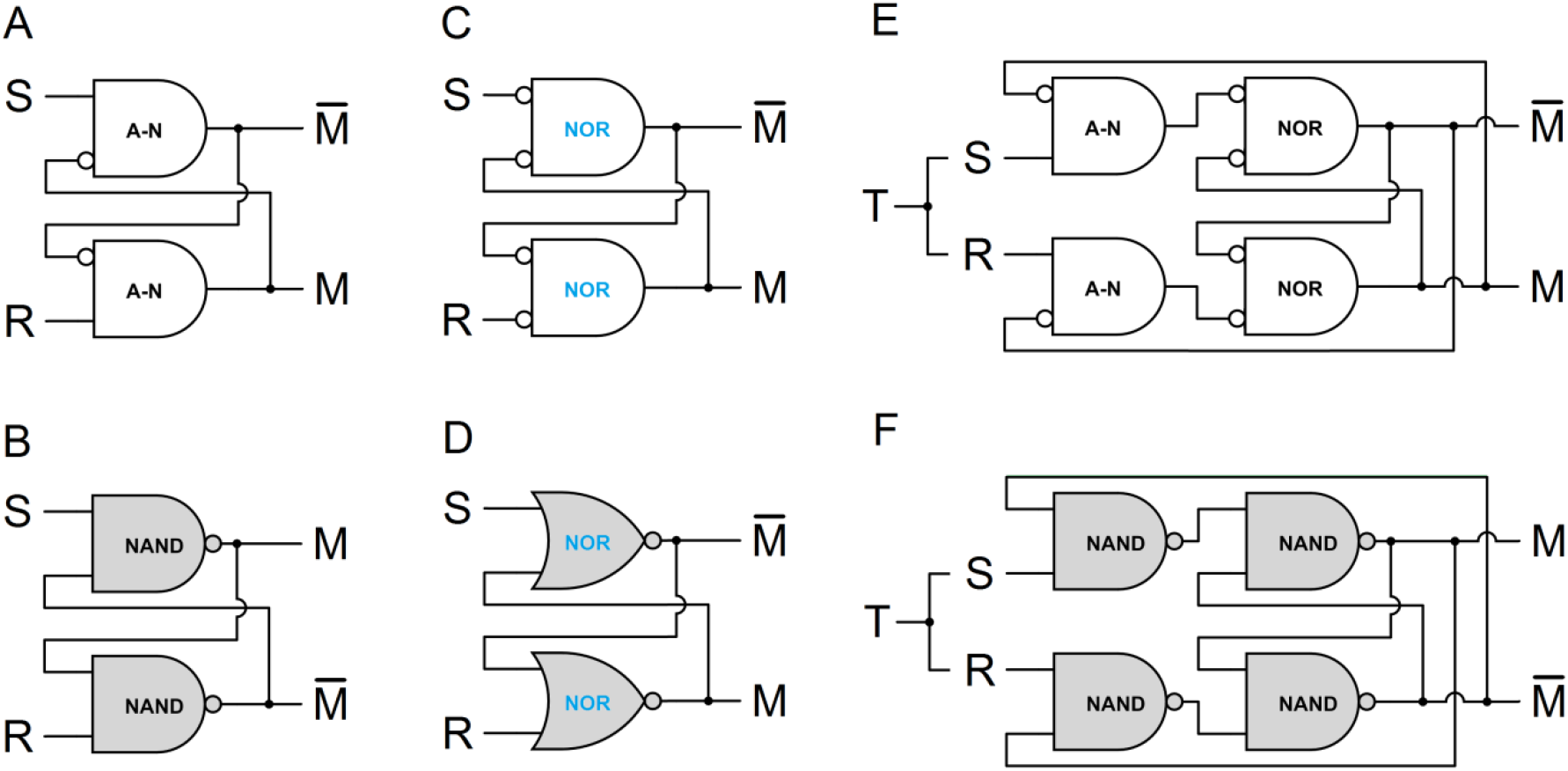
Neural and electronic flip-flops and oscillators. Each white network performs the same logic function as the gray network below it. The white networks are composed of logic gates from Fig 1 that can be implemented with neurons or with electronic components. The gray diagrams are widely known logic circuits and are composed of logic gates commonly used in electronic logic circuits. **A, B**. Active low set-reset flip-flops. **C, D**. Active high set-reset flip-flop composed of NOR gates. **E, F**. JK flip-flops or toggles.

Figs 2C and 2D show identical networks, illustrated with logically equivalent component symbols (Figs 1C and 1E). Each network in each of the other two pairs can be derived from the other network simply by moving each negation circle from one end of a connection to the other. If a circle is moved past a branch point to an output, the output is inverted. Moving the negation circles does not change the logic of the network, but it means each logic gate in the white networks can be implemented with a single neuron from Fig 1. Although this is a minor modification, the white networks that contain AND-NOT gates are likely new to engineering because the AND-NOT gate is virtually never used as a building block in logic circuit design.

Figs 2A and 2B show active low set-reset (SR) flip-flops. The S and R inputs are normally high. A brief low input S sets the memory M to 1, and a brief low input R resets it to 0. Negating the inputs of Fig 2A produces the active high SR flip-flop of Fig 2C. The S and R inputs are normally low. A brief high input S sets the memory M to 1, and a brief high input R resets it to 0. A problem with the SR flip-flops is that an error occurs if both S and R are low simultaneously for the active low flip-flops (Figs 2A and 2B), or simultaneously high for the active high flip-flops (Figs 2C and 2D).

Figs 2E and 2F show JK flip-flops. The advantage of the JK flip-flop is that if S and R are both high simultaneously, the flip-flop state is inverted because one of the inputs is suppressed by one of the outputs. This means the JK flip-flop can be configured as a toggle by linking the Set and Reset inputs, as illustrated in the figures with the single input T. A JK flip-flop configured in this way functions as a toggle only for high inputs of short duration. If the duration is too long, the outputs will oscillate.

##### 2.1.3.2. Flip-flop ring oscillators

An inverter is a logic operator that has a single input X and an output that is the opposite of X, i.e., NOT X. An odd number of three or more inverters connected sequentially in a ring produces periodic bursts as each gate inverts the next one. The odd number of inverters makes all of the inverter states unstable, so the states oscillate between high and low.

The flip-flop ring oscillator is a generalization of the inverter ring oscillator. It is composed of two or more simple flip-flops, with both outputs of each flip-flop connected as inputs to the next one. Fig 3 shows four flip-flop ring oscillators.

**Fig 3.**
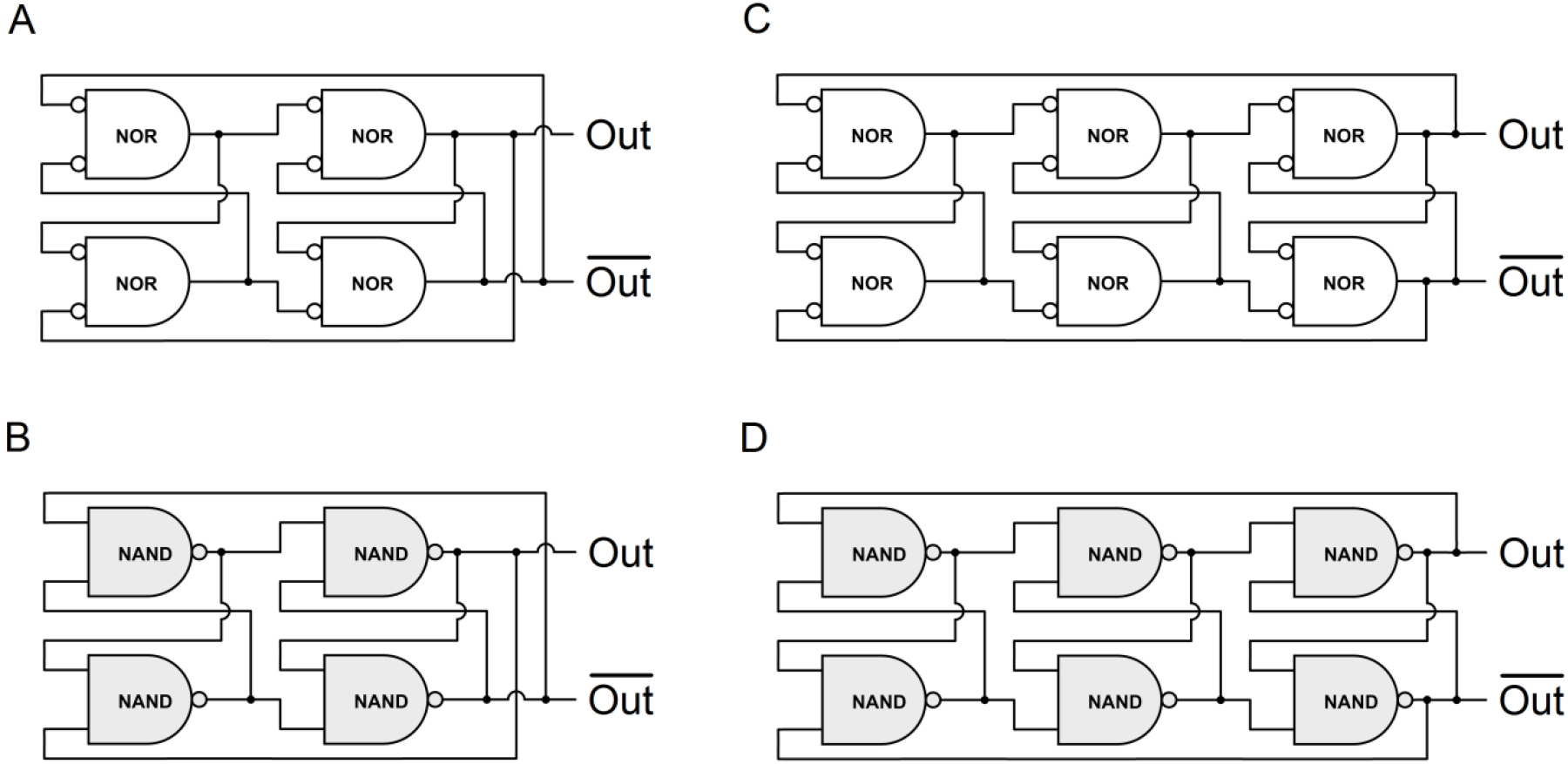
Flip-flop ring oscillators. The white and gray networks are composed of flip-flops from Figs 2C and 2B, respectively.

As in Fig 2, each white network in Fig 3 performs the same function as the gray network below it. Each network in each pair can be derived from the other by moving the negation circles from one end of a connection to the other. All of the gates (NOR and NAND) are commonly used in electronic logic circuits, but the NOR gate can be implemented by a neuron and is therefore white and represented by Fig 1C. The gray networks are light gray because they are composed of NAND gates that are common in electronic logic circuits, but unlike the gray networks in Fig 2, the flip-flop ring oscillator is apparently a new concept.

A flip-flop ring can have any number of flip-flops greater than one, but they must be connected differently depending on whether the number is even or odd. If each gate has input to the next gate ahead, as shown in Fig 3, but with no connection between the first and last flip-flop, all states are stable with alternating states horizontally and vertically. The first and last gates in each row have opposite states if the number of flip-flops is even and same states if the number is odd. The states are made unstable, as shown in Fig 3, by connecting opposite rows if the number of flip-flops is even and connecting to the same row if the number is odd. That is, the even number of flip-flops is connected like a Möbius strip, and the odd number like a ring.

### 2.2. Stomatogastric nervous system

#### 2.2.1. Schematic of the stomatogastric nervous system

A diagram of the stomatogastric nervous system is shown in Fig 4A, with simplified diagrams of the two CPGs in Figs 4B and 4C. Small triangles indicate excitatory synapses, and small circles indicate inhibitory synapses. Prominent features include mostly inhibitory synapses and pairs of cells with reciprocal, inhibitory input. Fig 4B shows three such pairs and Fig 4C has four. Several of the large circles in Figs 4B and 4C represent more than one cell, so most of the pairs represent multiple pairs in Fig 4A. Some of the cells have reciprocal, inhibitory inputs with more than one other cell.

**Fig 4.**
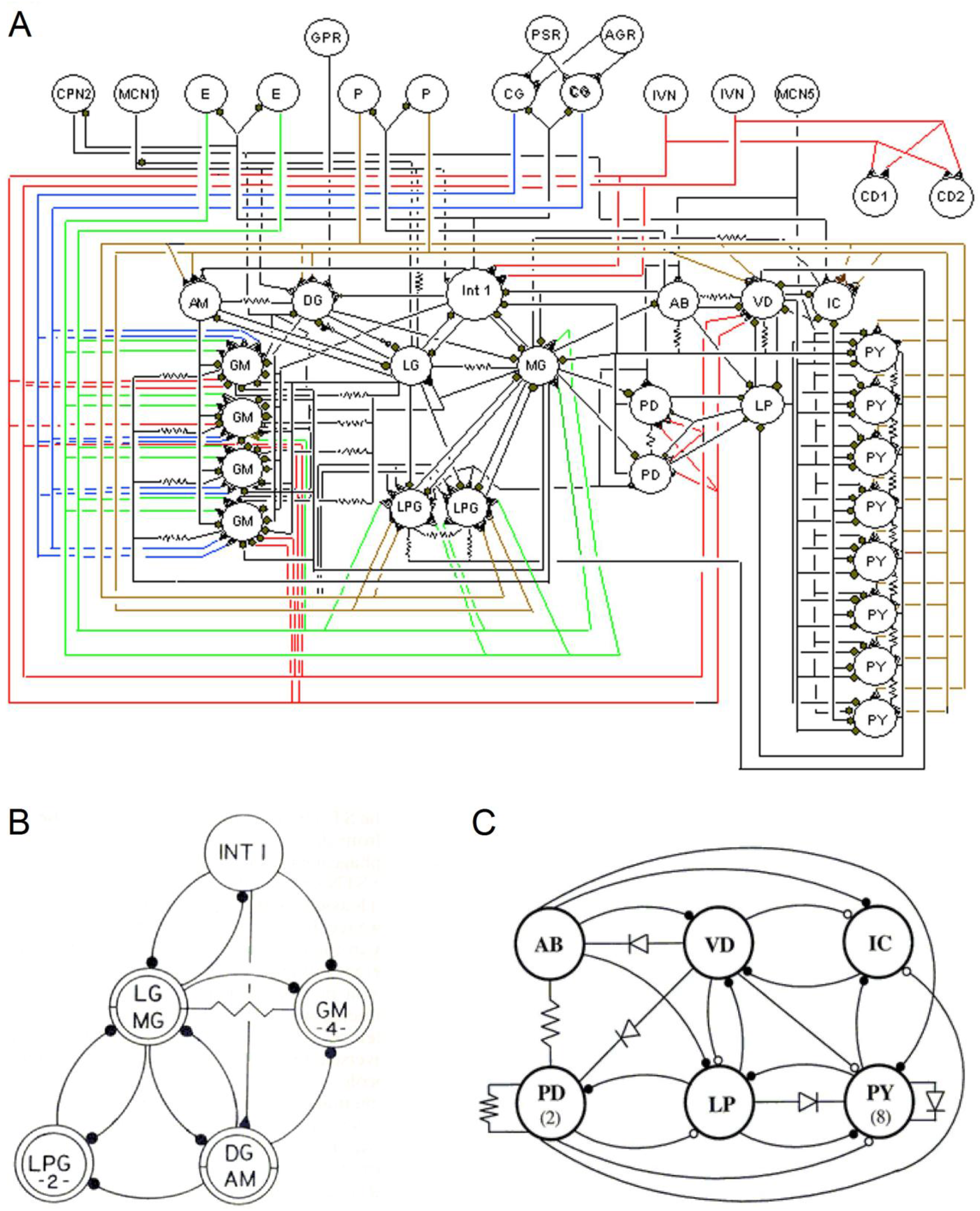
Stomatogastric nervous system. Triangles indicate excitatory synapses. Closed circles represent glutamatergic inhibitory synapses and open circles cholinergic inhibitory synapses. Resistors indicate electrical connections and diodes are rectifying connections. **A**. The stomatogastric nervous system. The gastric CPG is on the left and the pyloric CPG is on the right. At the top are excitatory neurons, brain cells, and sensory receptor cells. **B**. Simplified version of the gastric CPG. **C**. Simplified version of the pyloric CPG. (Diagrams courtesy of Allen Selverston [8].)

#### 2.2.2. Pyloric oscillations

Table 3 gives a brief description of the neurons of the pyloric CPG.

**Table 3.** Pyloric neurons. (Table courtesy of Allen Selverston [8].)

| # of copies | Name | Abbr. | Muscle | Transmitter |
| --- | --- | --- | --- | --- |
| 1 | anterior burster | (AB) | interneuron | glu |
| 2 | pyloric dilator | (PD) | cpv 1&2 | ach |
| 1 | lateral pyloric | (LP) | p1 | glu |
| 1 | ventricular dilator | (VD) | cv 2 | ach |
| 1 | inferior cardiac | (IC) | cv 3 | glu |
| 8 | pyloric | (PY) | p 2-4, 7-8, 10-11 | glu |

Fig 5 shows the results of 17 recordings of pyloric neuron bursts in American lobsters (*H. americanus*) [9]. (See Fig 4C.) As the authors point out, the bursts appear to have four distinct phases: 1) PD, 2) IC, 3) LP, and 4) VD, PY, and AM. The AM bars are light gray because that neuron was bursting in only 7 of the recordings.

**Fig 5.**
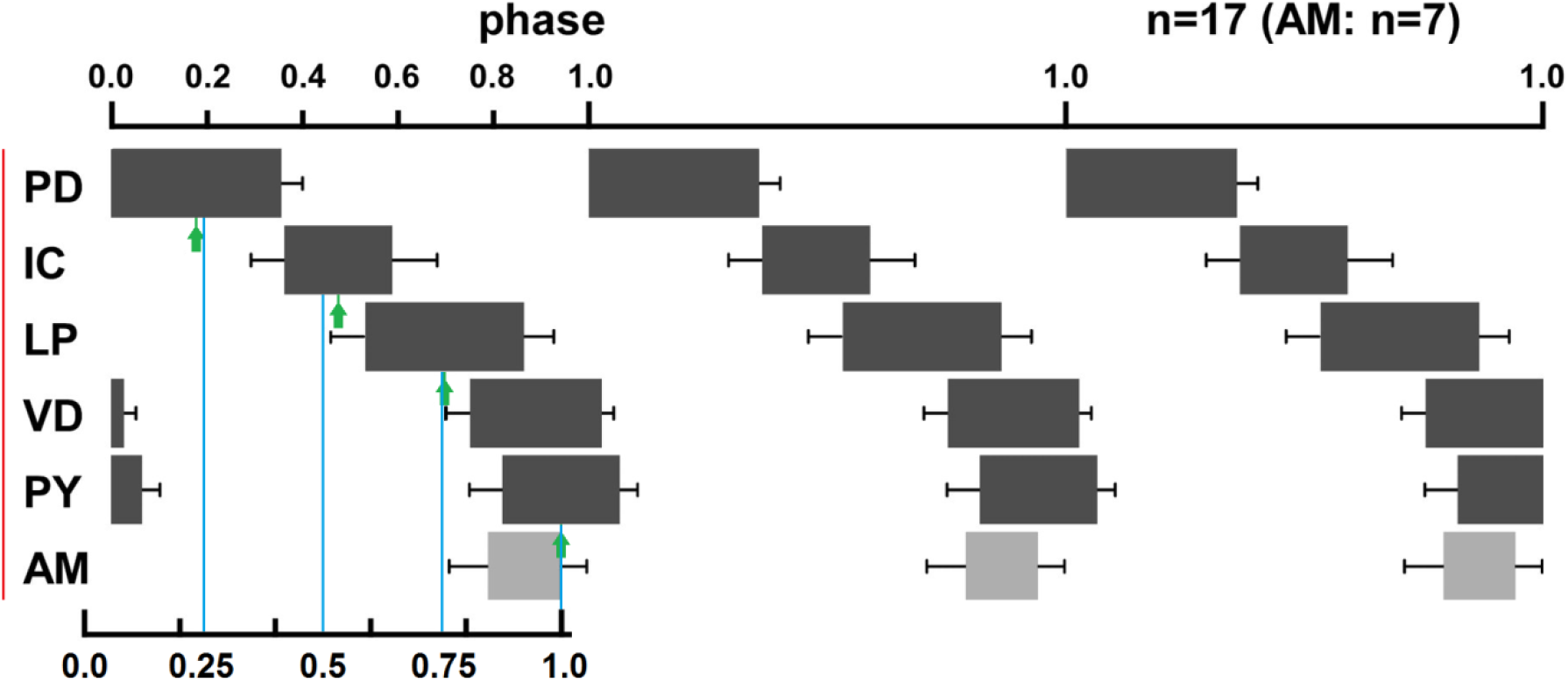
Pyloric phase and burst relationships. Three cycles of pyloric CPG bursts are shown. Error bars indicate standard deviations. (Adapted from [9].)

The colored parts and the copy of the period bar at the bottom of the figure were added to the original figure. The red line is used to align these graphs with the simulation graphs. The blue lines are uniformly distributed over the period (from pixel counts in the original image). The green arrows indicate the centers of four of the bursts (again from pixel counts). The arrows and blue lines show the burst centers are close to a uniform distribution over the period, especially considering the small sample size (17 recordings) and the magnitudes of the standard deviation bars.

The observation above - that the phases of four of the American lobster’s pyloric oscillations, as determined by the burst centers, are approximately uniformly distributed over the period - is apparently new. It will be seen in the simulations that different CPGs can produce oscillations with phases uniformly distributed at the burst centers, at the burst onsets, or both.

Table 4 shows the approximate burst durations of the oscillations in Fig 5 as a proportion of the period (from pixel counts in the original image).

**Table 4.** Pyloric burst durations in Fig 5.

| Neuron | Burst duration |
| --- | --- |
| PD | 0.36 |
| IC | 0.22 |
| LP | 0.33 |
| VD | 0.27 |
| PY | 0.25 |
| AM | 0.15 |

### 2.3. Comparing pyloric oscillations to simulations of model CPGs

#### 2.3.1. Classifying pyloric cells and comparisons with different models

The pyloric neurons considered here were chosen because they appear in Fig 5 and in both Tables 3 and 4, i.e., PD, LP, VD, IC, and PY. Categorizing a neuron as gastric or pyloric is complicated by the several connections between the two CPGs. Some neurons can fire with either gastric or pyloric rhythm, depending on neuromodulatory or sensory input [10-14]. Some neurons vary by species. The AM neuron, for example, fires mostly in gastric time in *C. borealis* and *P. interruptus* [12, 15]. It is either silent or fires in pyloric time in *H. Americanus* [9], and it fires in pyloric time in *H. gammarus* [16].

How well the model CPGs match the pyloric oscillations depends on the model’s oscillations’ burst durations and how the phases are distributed over the period. The pyloric neurons considered here produce short and long burst durations, averaging approximately 0.25 of the period for the IC, VD, PY cells and 0.35 for the LP and two PD cells (Table 4). The burst centers are approximately uniformly distributed over the period, as shown in Fig 5. Model CPGs based on the JK toggle produce oscillations that have burst durations of 1/4 and 3/8 (≈ 0.37) of the period, closely matching the pyloric burst durations. However, the burst onsets, not the centers, are uniformly distributed over the period. Consequently, the JK oscillators’ burst durations are a somewhat better match with the pyloric oscillations than the phases. Model CPGs based on flip-flop rings produce burst durations of 3/8 of the period, closely matching the pyloric LP and PD cells. Both the burst centers and burst onsets are uniformly distributed over the period. Consequently, the flip-flop rings’ phases match the pyloric somewhat better than the burst durations.

#### 2.3.2. Simulation and comparison methods

Since the model CPGs are illustrated with standard logic symbols, they can be constructed with ordinary electronic components or simulated with electronic circuit software. Two model CPGs were simulated in CircuitLab.

The CPG models that contain AND-NOT gates were simulated only in MS Excel. Since the AND-NOT gate is virtually never used in electronic circuit design, it normally is not found in simulation software. An AND-NOT gate could be composed of an AND gate and a NOT gate, but that would be simulated with two gate delay times. The electronic simulations are only for illustration. Because the neuron responses are the same as Boolean logic gates (Tables 1 and 2), all of the electronic implementations should match the neural implementations.

For the Excel simulations, the number t_i_ represents the time after i neuron delay times. The neurons’ outputs were initialized at time t_0_ = 0 as given in the text. For i > 0, the output of each neuron at time t_i_ was computed as a function of the inputs at time t_i-1_ according to Tables 1 and 2.

The model CPGs’ periods depend on the delay times of the component neurons. The delay times required for model CPGs to match the pyloric period were computed for each model. For comparing oscillations, the simulation graphs of each model CPG’s oscillations were stacked in a single image (as in Fig 5). The size of this image was adjusted to match the simulation period with the STG period in Fig 5. Because the centers of the pyloric bursts are distributed approximately uniformly over the period, the center of a burst bar of one of the pyloric oscillations from Fig 5 was aligned with the center of a burst of one of the simulated oscillations. The other pyloric oscillations of Fig 5 were then aligned vertically with the closest fits in the stack of simulated oscillations and horizontally with the red vertical line in Fig 5.

## 3. Results and Discussion

### 3.1. Simulation results

#### 3.1.1. Flip-flop ring oscillator

##### 3.1.1.1. The flip-flop ring oscillator implemented with neurons

Fig 6 shows a comparison of a flip-flop ring oscillator and the pyloric CPG’s four distinct phases.

**Fig 6.**
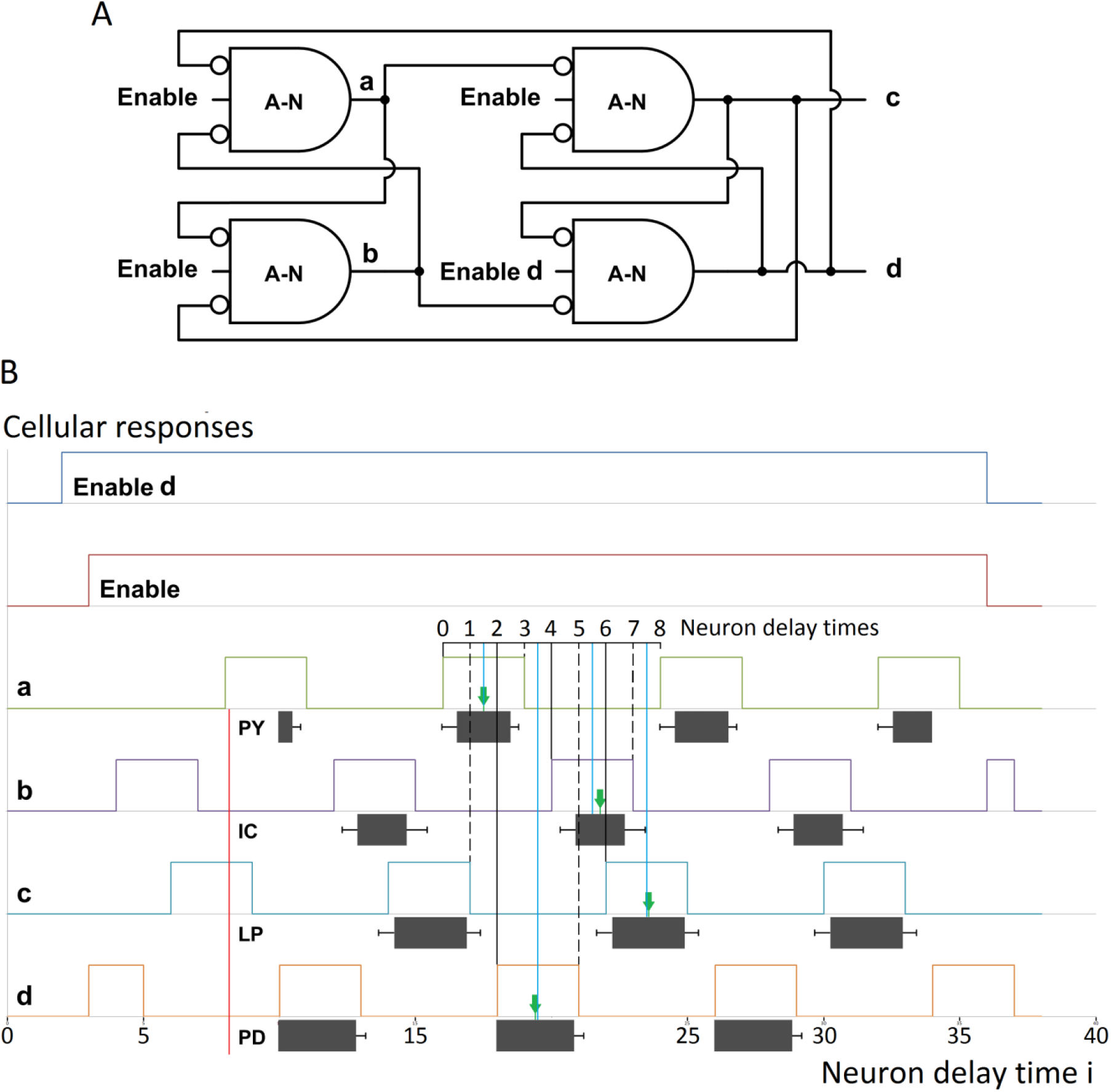
Neuron implementation of the flip-flop ring oscillator. **A**. The flip-flop ring oscillator of Fig 3A with excitatory enabling inputs from the brain or sensory cells. **B**. A comparison of a simulation of the flip-flop ring oscillator with four of the oscillations in Fig 5 that represent the pyloric CPG’s four distinct phases. The flip-flop ring’s four oscillations have the same period, eight neuron delay times. Their phases are uniformly distributed over the period at both the burst onsets, indicated by black lines, and at the burst centers, indicated by blue lines. The black lines and dashed lines together show that at each delay time, one of the four gates changes states. As in Fig 5, the green arrows indicate the centers of the pyloric bursts, and the close fit with the blue lines shows that the oscillator and pyloric phases, as determined by the burst centers, are approximately the same.

The flip-flop ring in Fig 6A is composed of AND-NOT gates from fig 1B instead of NOR gates of Fig 1C so the cells can be enabled by excitatory input as needed. When enabled, the AND-NOT gates function the same as NOR gates. The enabling input could be from the brain or sensory receptor cells, as described in the caption for Fig 4A. Fig 4A shows that nearly all STG cells have at least one excitatory input.

Fig 6B shows that a slightly different enabling input to cell d initializes its high output first to give the flip-flop ring an asymmetry for initialization. In the STG, neuromodulatory inputs are necessary to initiate the oscillation [8]. For the electronic simulation in the next section, the software initializes the states.

The flip-flop ring’s burst durations are three delay times, or 3/8 of the period. Table 4 shows the LP and PD burst durations of 0.33 and 0.36 are close to the flip-flop ring’s 3/8 ≈ 0.37. Only the IC, PY, and VD bursts (0.22, 0.25, 0.27) are substantially shorter than the flip-flop ring’s 0.37, although Fig 6B shows the flip-flop ring’s burst durations are close to the extents of the pyloric IC and PY standard deviation bars. The pyloric period is 1.35 ± 0.18 seconds [9]. For the flip-flop ring’s eight neuron delay times per cycle, matching the pyloric period would mean an average neuron delay time of about 170 ms.

The alternative flip-flop ring composed of NAND gates (Fig 3B) has two problems with respect to modeling the STG. It is not clear that a single neuron can function as a Boolean NAND gate. That would require a neuron that is active when either of two inhibitory inputs is low, rather than both. Second, the NAND ring oscillator gives the same uniformly distributed phases as the ring with NOR gates, but the pulse and quiescent durations are reversed, i.e., long pulses (5/8) and short quiescent intervals (3/8). These durations are quite far from the pyloric. Both of these problems are consistent with the hypothesis that the STG network architecture is similar to the NOR flip-flop ring of Fig 3A.

##### 3.1.1.2. The flip-flop ring oscillator implemented with electronic components

To verify the operation of the flip-flop ring oscillator, Fig 7 shows the same comparison as Fig 6, with a CircuitLab simulation of an electronic implementation of the flip-flop ring of Fig 3A instead of the neural implementation. The result in Fig 7B is virtually identical to Fig 6B.

**Fig 7.**
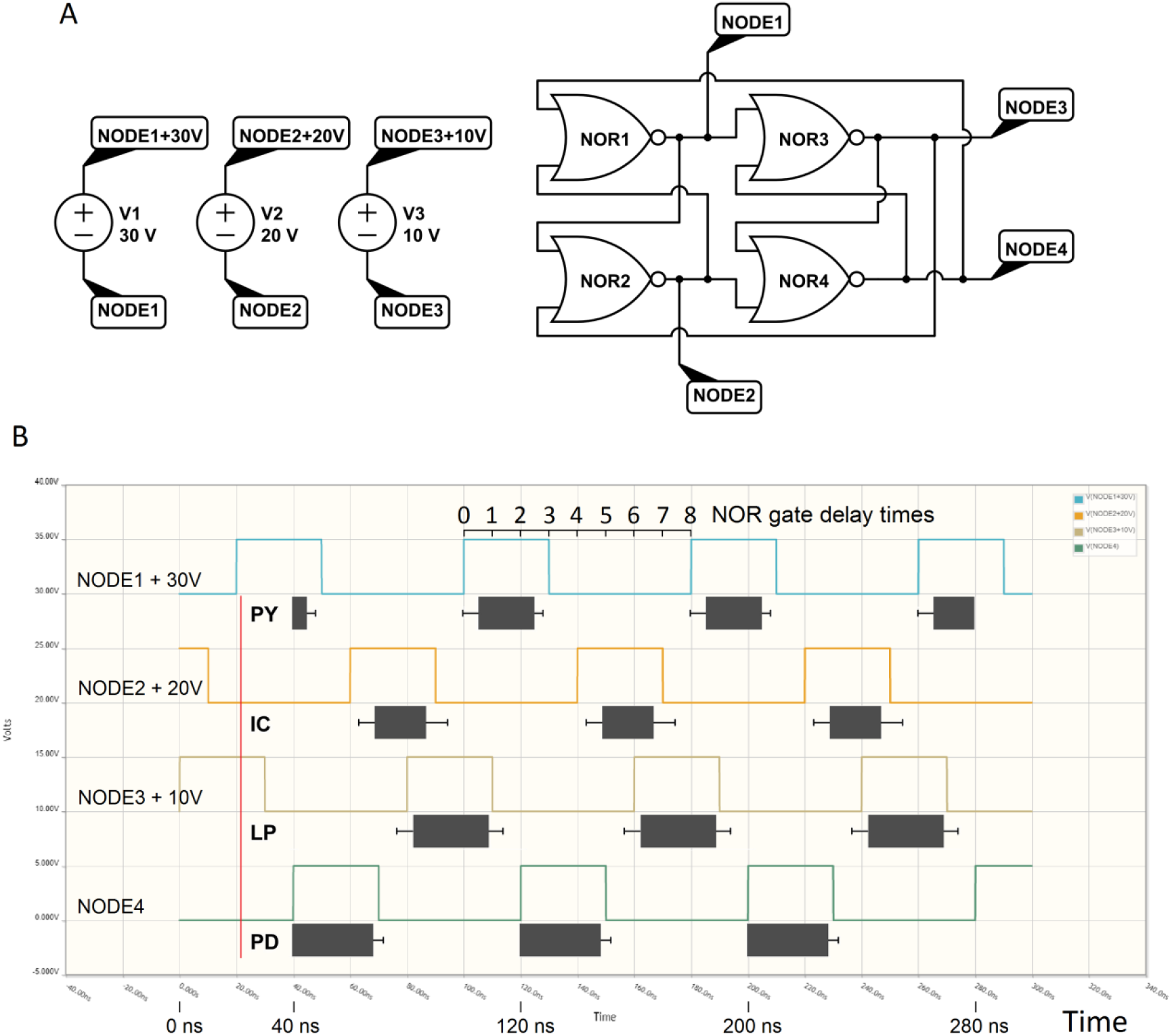
Electronic implementation of the flip-flop ring oscillator. **A**. Flip-flop ring of Fig 3A (with NOR gate symbols of Fig 1E). **B**. CircuitLab simulation of the flip-flop ring with the pyloric oscillations of Fig 5.

#### 3.1.1.3. An extension of the flip-flop ring oscillator

An extension of the flip-flop ring oscillator can model six of the pyloric neurons, including the second PD neuron and VD. This extended ring oscillator also shows that a NOR gate (neural or electronic) can have reciprocal, inhibitory inputs with more than one other gate. Fig 4 shows the STG has several such neurons (e.g., the pyloric LP and VD). Although pairs of NOR or NAND gates with reciprocal, inhibitory inputs are quite common in electronic logic circuits, electronic gates apparently do not have reciprocal, inhibitory inputs with more than one other gate.

Figs 8A and 8B show an extension of the flip-flop ring oscillator in Fig 3A. The two networks are composed of logically equivalent NOR gates from Figs 1C and 1E, respectively. The general NOR gate in Fig 1D with three inputs is also used. Since the networks in Figs 8A and 8B are logically equivalent, for convenience an electronic simulation of Fig 8B is shown in Fig 8C.

**Fig 8.**
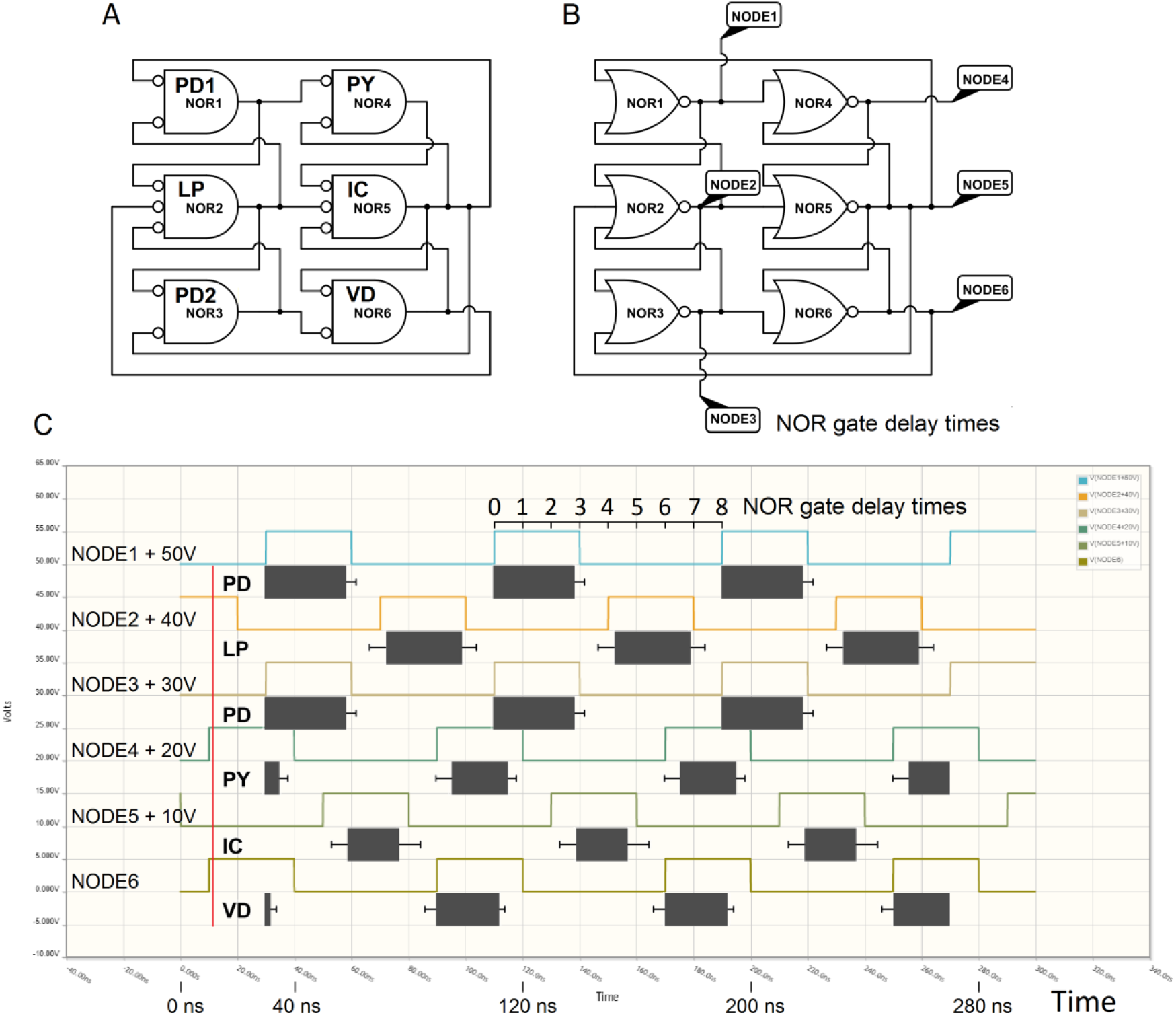
An extended flip-flop ring oscillator. This extension of the flip-flop ring oscillator of Fig 3A models six of the pyloric neurons, including the second PD neuron and VD. It also demonstrates that gates can have reciprocal, inhibitory inputs with more than one other gate. **A**. An extension of the flip-flop ring oscillator. **B**. The same network as in Fig A, illustrated with the most commonly use NOR gate symbol (Fig 1E). **C**. A CircuitLab simulation of the model CPG in Fig B. Like the flip-flop ring simulations in Figs 6 and 7, the phases are uniformly distributed at the burst centers, and the burst durations are 3/8 of the period.

The CircuitLab software assumes a 10 nanoseconds delay for each gate. The simulations in Figs 7B and 8C show that the flip-flop ring oscillations have a period of 80 ns. The pyloric oscillations of the American lobster have a period of 1.35 ± 0.18 sec [9]. So the electronic and lobster frequencies differ by a factor of about 17 million.

#### 3.1.2. JK toggle

##### 3.1.2.1. Two configurations of the JK toggle as oscillators

Fig 9A shows the JK NFF of Fig 2E configured as a toggle. The simulations in Figs 9B and 9C show the toggle’s output with an oscillating input and with a continuously high input.

**Fig 9.**
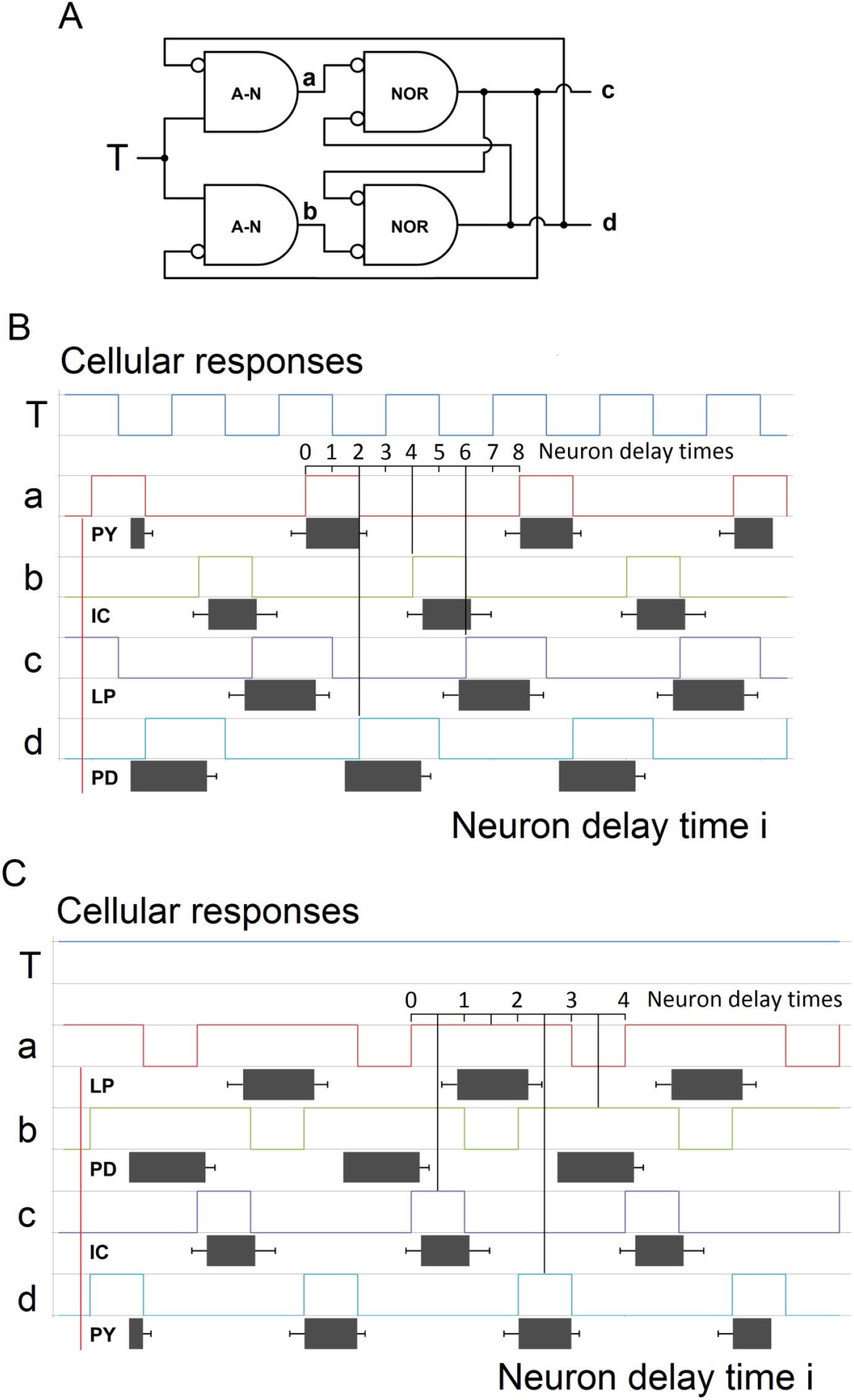
JK toggle configured as an oscillator. **A**. The JK toggle of Fig 2E. **B**. Simulation of the toggle with input from an oscillator. **C**. Simulation of the toggle with continuously high input.

Fig 9B shows the simulated JK toggle with an oscillating input, possibly from an intrinsically bursting neuron or pacemaker neuron. The simulation shows two oscillations with burst durations of 0.25, nearly a perfect match with the pyloric PY and IC bursts of 0.25 and 0.23 (Table 4). This is also a close match with the pyloric VD burst of 0.27 (not shown in the simulation). The other two burst durations are 3/8 ≈ 0.37 like the flip-flop ring oscillator of Fig 6, a reasonably close match with the pyloric LP and PD bursts of 0.33 and 0.36. However, the JK toggle’s phases are uniformly distributed at the burst onsets, as indicated by the black lines, not at the burst centers. So the phases do not match the pyloric CPG quite as well as the flip-flop ring oscillator of Fig 6.

Like the flip-flop ring (Figs 6-8), the JK toggle’s period is eight neuron delay times. So to match the pyloric period of 1.35 sec, the average neuron delay time must be about 170 ms. Since the JK toggle is driven by an oscillating input, the CPG’s period also depends on the period of the input. For a JK toggle with oscillating input to oscillate successfully, the burst duration of the input must be about two neuron delay times. For the JK toggle to have burst durations of 1/4 and 3/8 of the period, the quiescent interval of the input must also be two neuron delay times. That makes the period of the oscillating input about 0.68 sec, or a frequency of about 1.47 Hz.

Fig 9C shows a simulation of the JK toggle with a continuously high input. Like the toggle with oscillating input in Fig 9B, this network has two oscillations with burst durations of 0.25 that closely match the pyloric PY and IC bursts of 0.25 and 0.23. The phases are uniformly distributed at the burst centers, indicated by the black vertical lines, so all of the phases also show a good fit. However, the JK toggle’s other two burst durations are 0.75, substantially greater than the pyloric burst durations of 0.33 and 0.36. The model CPG’s period is the sum of four neuron delay times. For this period to match the pyloric period, the average neuron delay would be about 0.34 sec.

The pyloric PY was placed at the bottom to match the short burst of oscillation d. However, the order of the pyloric oscillations is not the reverse of the order in all of the previous simulations. The order is different because the order of the simulated oscillations is actually different, not simply reversed.

##### 3.1.2.2. An extension of the JK toggle

Fig 10 shows that an extension of the JK toggle produces a close match with six of the pyloric oscillations. Like the extension of the flip-flop ring in Fig 8, a cell has reciprocal, inhibitory inputs with more than one cell. Also like the JK toggle, the extended version’s oscillations are uniformly distributed by burst onsets, indicated by vertical lines, so not all of the phases match the pyloric phases closely.

**Fig 10.**
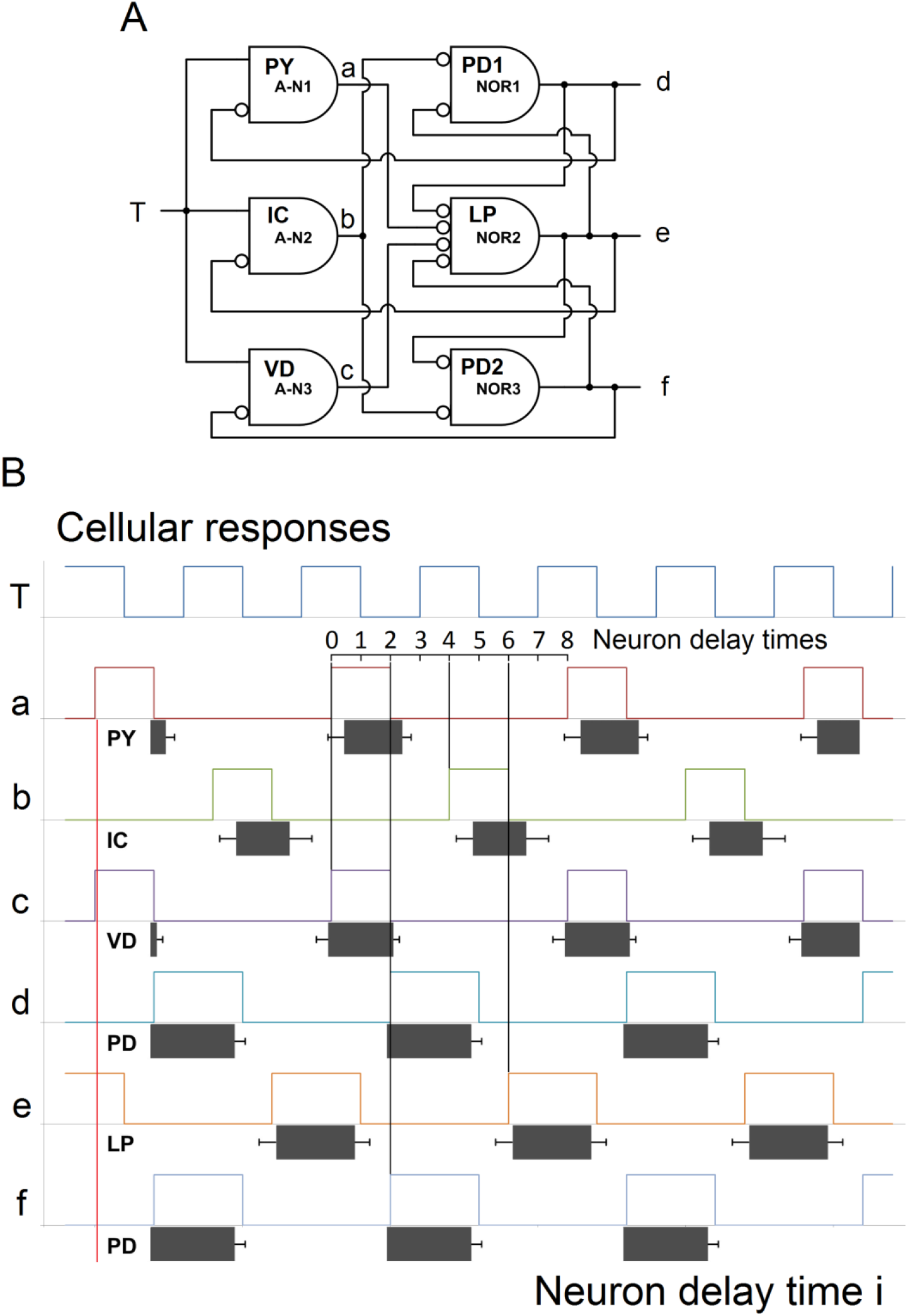
An extended JK toggle.

##### 3.1.2.3. The problem of modeling a mechanism with several attributes

Finally, Fig 11 illustrates the difficulty in designing a CPG that generates an arbitrary set of oscillations with various phases and burst durations. Fig 11A shows an extension of the JK toggle in Fig 9A that is different from the extension in Fig 10A and similar to the extension of the flip-flop ring in Fig 8A.

**Fig 11.**
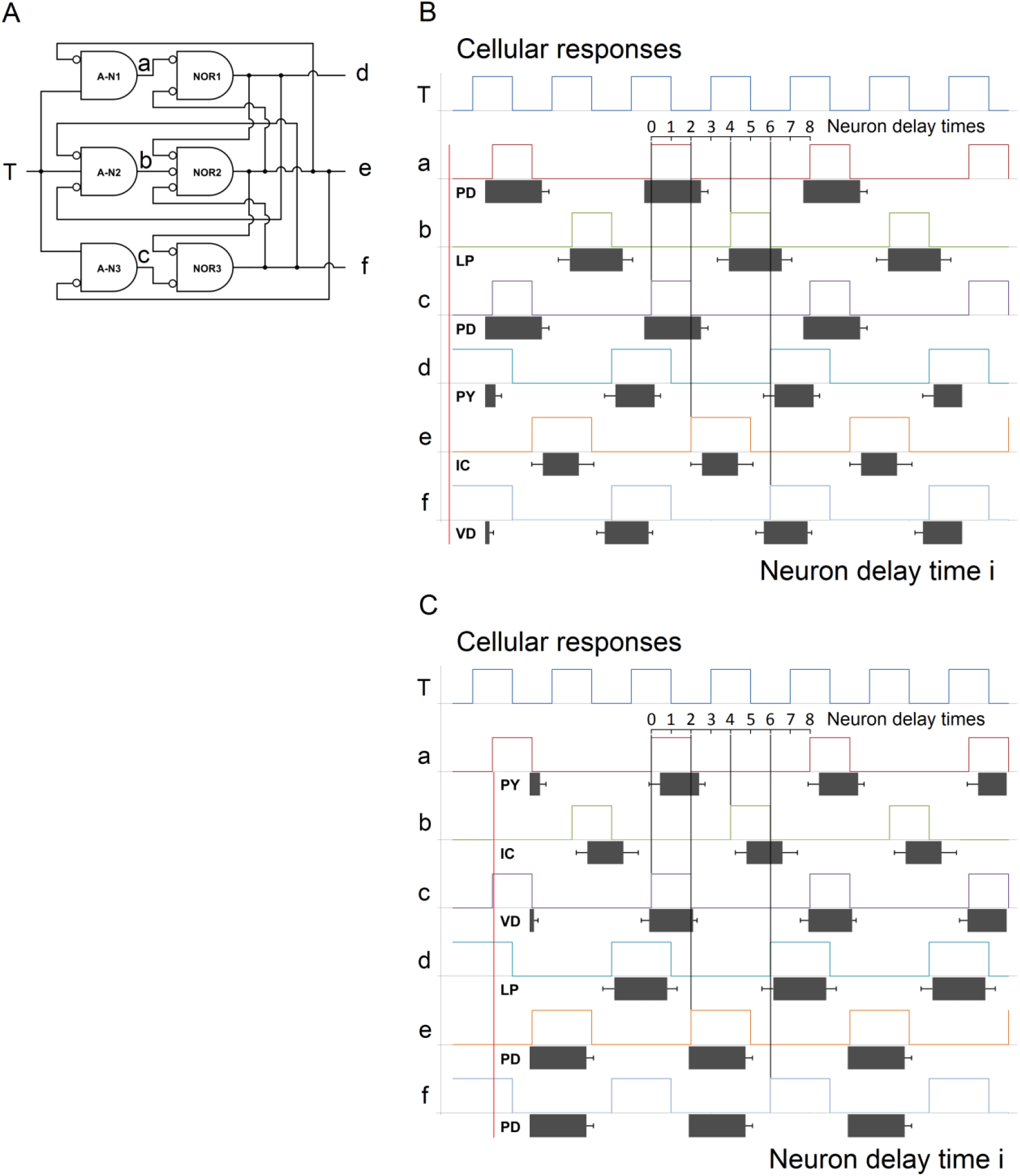
A different extension of the JK toggle. In spite of basic similarities between the oscillations of the model CPG and the pyloric CPG, they do not come close to matching.

The model CPG in Fig 11A has all of the similarities to the pyloric CPG that are listed in the abstract, and more: Like Fig 10B, the simulation in Figs 11B and 11C shows three oscillations with bursts of 1/4 of the period and three with bursts of 3/8. A matched pair of oscillations at each burst duration is 180 degrees out of phase with the third oscillation of that burst duration.

Yet the oscillations do not match even approximately. If the oscillations are aligned by the best fit of phases, as in Fig 11B, the burst durations are reversed. If the oscillations are aligned by the best fit of burst durations, as in Fig 11C, at least one pair of oscillations is out of phase by about 180 degrees.

### 3.2. Predictions

#### 3.2.1. Previous predictions supported by STGs

Several STG phenomena support NFF predictions in a previous paper [4]. There the NFF output M or 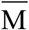 was shown to produce seven known characteristics associated with short-term memory formation, retention, retrieval, termination, and errors. It was also shown that M and 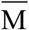 together predict eight unknown phenomena. STGs verify five of the predictions: The two output neurons were predicted to have 1) close proximity; 2) reciprocal, 3) inhibitory inputs; and 4) complementary outputs. When the neurons change states, 5) the neuron with high output changes first. Predictions 1-3 can be seen here in the NFFs in Fig 2, and predictions 4 and 5 can be seen in the outputs c and d in the two simulations of the JK NFF in Fig 9.

At least three pyloric neuron pairs satisfy these predictions: PD-LP (two pairs) and IC-VD (one pair). The STG consists of about 30 neurons, so the neurons are in close proximity. Figs 4A and 4C show each pair has reciprocal, inhibitory inputs. Fig 5 shows each pair has complementary outputs, and that when each pair changes states, the neuron with high output changes first.

#### 3.2.2. Testable predictions of constructed neural networks

Any of the networks in the figures could be constructed with actual neurons and tested for their predicted behavior. (Constructing and testing NFFs were also discussed in [4, 5].) Fig 12 shows a few simple possibilities.

**Fig 12.**
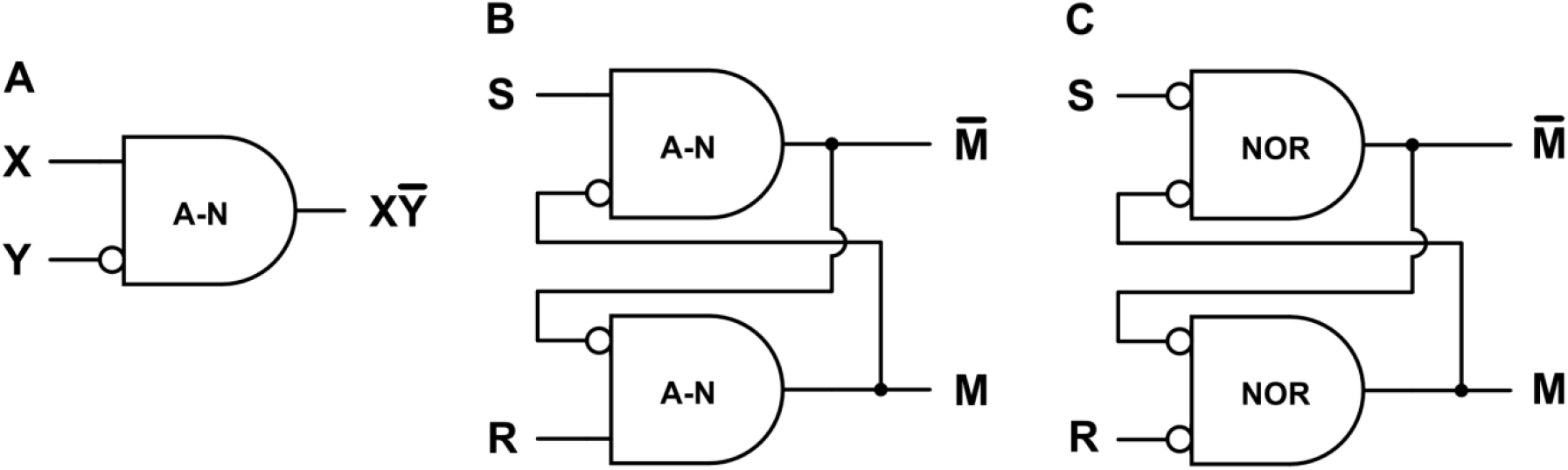
Simple neural networks that can be constructed and tested for predicted behavior. The networks are from Figs 1 and 2.

The predicted behavior Fig 12A is indicated by Table 1. For the NFFs in Figs 12B and 12C, brief input from S sets M high and brief input from R resets M low. Recall the NFF in Fig 12B is active low, i.e., the inputs are normally high and brief low inputs from S and R invert the state. Fig 12C requires spontaneously active neurons.

## 4. Acknowledgements

Simulations were done in CircuitLab and Excel. Figures were created in CircuitLab and MS Paint. The author would like to thank Duncan Watson, Arturo Tozzi, Robert Barfield, David Garmire, Paul Higashi, Anna Yoder Higashi, Sheila Yoder, and especially Ernest Greene and David Burress for their support and many helpful comments.

## Notes

### Competing Interest Statement

The authors have declared no competing interest.

### Summary of Updates

The abstract and introduction were rewritten, and the section headings were changed to be consistent with the standard Introduction, Methods, Results and Discussion.

